# Nondestructive Detection and Quantification of Dysprosium in Plant Tissues

**DOI:** 10.1101/2025.01.07.631589

**Authors:** Edmaritz Hernández-Pagán, Kanjana Laosuntisuk, Alex T. Harris, Allison N. Haynes, David Buitrago, Cyprian Rajabu, Michael W. Kudenov, Colleen J. Doherty

**Affiliations:** Department of Molecular and Structural Biochemistry, North Carolina State University, Raleigh, NC; Department of Electrical and Computer Engineering, North Carolina State University, Raleigh, NC

**Keywords:** Phytomining, Rare Earth Elements, Dysprosium Detection, Spectral Imaging, Agromining

## Abstract

**Background:** The growing demand for rare-earth elements (REEs), particularly dysprosium (Dy), in part driven by clean energy technologies, underscores the need for sustainable extraction methods. Recovery of Dy, particularly from geographically distributed waste sources is challenging. This gap positions phytomining—a technique using plants to accumulate metals— as a promising alternative. However, plant species differ in their ability to accumulate metals in high concentrations, necessitating efficient screening methods. In this study, we developed a high-throughput fluorescence-based assay to detect and quantify Dy uptake in plant tissues.

**Results:** Our Dy detection method exploits Dy’s unique spectroscopic properties for sensitive and efficient analysis, enabling detection of concentrations as low as 0.3 µM. By incorporating sodium tungstate (Na WO) as a fluorescence enhancer, we achieved robust emissions at 480 and 580 nm, facilitating Dy quantification in complex plant matrices. Additionally, time-resolved fluorescence techniques reduced background autofluorescence from plant tissues, enhancing signal specificity. Validation against Inductively Coupled Plasma Mass Spectrometry (ICP-MS) demonstrated strong correlation. Greenhouse trials confirmed the method’s utility for screening Dy accumulation in living plants and highlight the potential for rapid standoff detection.

**Conclusions:** This fluorescence-based approach offers a scalable, efficient tool for identifying Dy-accumulating plants, advancing phytomining as a sustainable strategy for REE recovery.

## Background

Rare-earth elements (REEs) comprise 17 elements, including scandium (Sc), yttrium (Y), and the lanthanides from lanthanum (La) to lutetium (Lu) [1]. The rising use of clean energy technologies has sharply increased the demand for REEs, particularly dysprosium (Dy) [2–4]. Dy is added to neodymium–iron–boron (NdFeB) permanent magnets to improve high-temperature resistance [5]. These magnets are used in hybrid electric vehicles, wind turbines, computer hard drives, home appliances, and other applications [2].

Extracting and processing REEs is challenging despite their relative abundance in the earth’s crust. REEs are usually dispersed and mixed with non-REE metals, making purification costly and potentially environmentally harmful [6]. China holds 37% of the global REE reserves and 87% of the global Dy reserves [3]. This spatial distribution of REEs has heightened interest in alternative sources to support the green energy transition [7,8].

Concerns over supply risks and environmental impacts have driven efforts to find new REE sources and reduce the carbon footprint of traditional mining [6,9]. Secondary sources like coal fly ash, acid mine drainage, and recycled magnets are being explored to ensure a sustainable REE supply [10,11].

Phytomining - using plants to extract metals from soil [12,13] - shows promise for sustainable Dy recovery. Some plants can tolerate, uptake, concentrate, and translocate metals to aerial tissue at levels significantly higher than, or even toxic to, other plants in the same environment [14,15]. Metal accumulation in plants varies considerably from species to species, but some plants can accumulate metal concentrations from 100 to 10,000 mg/kg or higher. Plants that accumulate metals to these high levels tend to be specific to each metal element [15]. Due to their ability to concentrate metals, such plants are valuable for extracting precious elements and cleaning contaminated soils [16]. Current nickel phytomining efforts have successfully identified nickel-accumulating species like *Pycnandra acuminata* and *Phyllanthus balgooyi,* which are being scaled to mine nickel commercially [17–20].

Efforts are ongoing to identify plant species suitable for REE phytomining. For example, plants in the *Phytolacca* genus are candidates for phytoming and phytoremediation [12,21,22]. *Phytolacca americana* can accumulate up to 1,040 mg/kg REEs in the leaf tissues when grown in REE-contaminated soils [23]. Expanding the knowledge of these and other species across various environments is crucial for developing effective REE phytomining methods.

To develop a successful Dy phytomining system for secondary REE sources, it is essential to rapidly identify plants with the capacity to accumulate high Dy levels in their leaf tissues. Conventional methods for measuring Dy in plants rely on Inductively Coupled Plasma-based techniques (ICP) such as ICP mass spectrometry (ICP-MS), ICP optical emission spectroscopy (ICP-OES), and ICP atomic emission spectroscopy (ICP-AES). ICP-MS can detect elements at concentrations down to parts per trillion (ppt), equivalent to 0.001 μg/L, and down to picomolar concentrations - 6.15 x10^-6^ μM Dy. In contrast, ICP-OES and ICP-AES can detect elements down to parts per billion (ppb), equivalent to 1 μg/L and down to nanomolar concentrations [24]. However, these techniques can require high amounts of plant tissue, and the sample preparation results in tissue destruction due to an acid digestion step, limiting single-plant and time course Dy detection and measurement [25]. Since plant species vary in REE uptake, developing non-destructive screening methods will help identify plants with the highest REE accumulation potential. Alternative measurement techniques include X-ray fluorescence (XRF). XRF microscopy was used to detect and map La and cerium (Ce) in *Dicranopteris linearis* growing on REE mine tailings in Ganzhou, Jiangxi Province, China. The results showed that La and Ce were colocalized in the pinnae and pinnules and were enriched in areas of dead tissue and leaf veins [26]. Similarly, XRF spectroscopy was implemented to identify Yttrium (Y) hyperaccumulating plants from herbarium specimens [27]. This screen identified plants previously described as high REE accumulating species, including *D. linearis* and *Blechnopsis orientalis*. In addition, this study identified 15 new REE-accumulating species. However, *P. americana,* a species known to accumulate REEs and preferentially accumulate heavy REEs, such as Dy, in the leaves [22,23], was not identified in the herbarium screen. Herbarium collections include plants from various locations. Therefore, the *P. americana* plants in the herbarium collection may have been collected from areas with low or no REEs in the soil. In addition to screening herbarium samples, Goudard *et al.* also employed XRF microscopy to examine the distribution of five other REEs in addition to Y (Ce, La, Nd, Praseodymium (Pr), and Ytterbium (Yb)) in *D. linearis* tissue. Y showed different distribution patterns compared to the other elements tested. These findings highlight XRF’s potential for measuring the concentration and distribution of REEs in plants.

Here, we present a method for screening Dy uptake in plants based on fluorescence. REEs’ spectroscopic properties allow high-throughput plant detection from limited materials. Lanthanide 4f-4f transitions produce multicolor emission from the ultraviolet-visible-near-infrared (UV-Vis-NIR) range, long luminescence decay, and narrow absorption and emission bands [28,29]. Dy’s emits yellow and blue under UV excitation [30]. Here, we describe a 96-well plate UV fluorescence assay for phytomining-relevant Dy concentrations, demonstrate its application to plant tissue, and compare it with ICP-MS. This high-throughput approach can screen plant tissues for Dy uptake using 96 well-plates, and we demonstrate its potential for standoff detection in in-field measurements.

## Methods

### 1. Dy absorbance and fluorescence measurement from DyCl_3_ solution to determine Dy excitation wavelength

Solutions containing 100 mM (16,250 mg/L) of Dy were prepared with DyCl_3_ in water at room temperature in a 96-well plate and agitated for 2 hours and 30 minutes. Absorbance measurements (BioTek Synergy H1 Multimode Reader, BioTek, USA) were taken from 250 - 700 nm at 10 nm intervals, from 250 - 700 nm at 10 nm intervals, and from 250 - 500 nm at 1 nm intervals. The wavelength of maximum absorbance obtained (350 nm) was used to excite the Dy ions in the solutions, implementing 60 μs time-resolved fluorescence and a 300 µsec delay collection time. The sensitizer, Na_2_WO_4,_ was added 0 and 0.8 μM Dy solutions for a final concentration of 0.25 M, and excitation wavelengths were tested to determine the highest Dy emission using 60 μs time-resolved fluorescence and a 300 µsec delay collection time. Emission was measured at 480 and 580 nm. The instrument’s general settings are as follows: the optics are on the top of the plate, the light source is a xenon flash, and the lamp energy is high.

### 2. General settings for Dy fluorescence assay in the plate reader

Dy emissions were measured using 60 µs time-resolved fluorescence and a 300 µs delay collection time. The excitation wavelength used was 275 nm, and the emission was collected from 450 to 700 nm. The instrument’s general settings are as follows: the optics are on the top of the plate, the light source is a xenon flash, and the lamp energy is high.

### 3. Dy fluorescence measurement from DyCl_3_ solution

In the 96-well plates, 10 µL of DyCl_3_ solutions were mixed with 100 μL Na_2_WO_4_ for a final concentration of 0.25 M. The concentrations of DyCl_3_ in this test were 0, 0.07, 0.1, 0.3, 0.5, 0.7, 1 µM. The concentrations of Dy in µg/L were as follows: 0, 0, 11.375, 16.25, 48.75, 81.25, 113.75, and 162.5. The plates were agitated at room temperature for 2 hours and 30 minutes before the fluorescence assay. Dy emissions were measured using time-resolved fluorescence as described above.

### 4. Preparation of Dy-spiked samples

*P. acinosa* aerial tissue was dried in an oven overnight at 90°C and 0.31 g of dried tissue was mixed with 31 mL of DyCl_3_ solution in Digitubes (SCP Science) to get final concentrations of 0, 0.005, 0.05, 0.02, 0.8, 3.2, and 25.6 µM (0, 0.8125, 8.125, 32.5, 130, and 4160 ug/L) of DyCl_3_ in the samples. The samples were used for Dy emission spectra measurements and ICP-MS. Dy emissions were measured using time-resolved fluorescence as described above.

### 5. *P. acinosa* growth and treatment with DyCl_3_ solution

*P. acinosa* seeds were stratified in concentrated sulfuric acid for 5 minutes with agitation and rinsed with DI water five times. Seeds were then surface sterilized with 50% bleach supplemented with 0.02% (v/v) Triton-X for 7 minutes with agitation and rinsed with DI water three times. Afterward, seeds were kept in sterile DI water in the dark at 25°C for two days and exposed to 12/12 h light/dark cycles for 7-10 days for germination. Germinating seeds were transferred to growing medium (OASIS^®^ ROOTCUBES^®^, Oasis grower solution, USA) containing 0.1x Hoagland solution (0.5 mM KNO_3_, 0.5 mM Ca(NO_3_)_2_·4H_2_O, 0.6% Sprint^®^138 chelated iron, 0.2 mM MgSO_4_·7H_2_O, 0.1 mM NH_4_NO_3_, 0.2 mM KH_2_PO_4_, 5 µM H_3_BO_3_, 1 µM MnCl_2_·4H_2_O, 0.08 µM ZnSO_4_·7H_2_O, 0.03 µM CuSO_4_·5H_2_O, 0.01 µM H_3_MoO_4_·H_2_O). Once the plants had four fully expanded leaves, plants were transferred to 0.25x Hoagland solution (1.25 mM KNO_3_, 1.25 mM Ca(NO_3_)_2_·4H_2_O, 1.5% Sprint^®^138 chelated iron, 0.5 mM MgSO_4_·7H_2_O, 0.25 mM NH_4_NO_3_, 12 µM H_3_BO_3_, 2 µM MnCl_2_·4H_2_O, 0.2 µM ZnSO_4_·7H_2_O, 0.08 µM CuSO_4_·5H_2_O, 0.02 µM H_3_MoO_4_·H_2_O) supplemented with 0, 5, 10, 15, and 20 mM DyCl_3_ solution (equivalent to 0, 812.5, 1,625, 2,437.5, and 3,250 mg/L of Dy). KH_2_PO_4_ was excluded from the Hoagland solution to prevent Dy precipitation with phosphate [22]. The leaf discs were cut from two areas of one leaf, leaf lamina and midrib, after growing in DyCl_3_ for seven days using a 2-mm diameter cork borer and placed in a 96-well plate. The discs were then submerged in 100 µM of 0.5 M Na_2_WO_4_, and the plates were agitated at room temperature for 2 hours and 30 minutes before taking a measurement. Dy emissions were measured using time-resolved fluorescence as described above.

### 6. Dy screening in intact plant tissue

A commercial deep UV Raman and fluorescence spectrometer by Photon Systems Inc. (DUV Raman/PL 200) was used to measure Dy signal from *P. acinosa* plants treated with a water solution containing 0, 10, and 50 mM (0, 1,625, and 8,125 mg/L) of DyCl_3_ following the fluorescence spectroscopy methods in [31]. The laser had a pulse frequency rate of 40 pulses per second, and 10 pulses with an energy of 5.2 μj were used with a diameter between 100-200 microns. After plant treatment for 48 hours, fluorescence emission spectra in the youngest leaves of the plants were measured using a 248 nm excitation wavelength. The emission at 480 nm and 575 nm wavelengths was used as the Dy signal. The stems and all the leaves of the plants were collected, pooled, and processed for ICP-MS measurements.

### 7. Dy analysis by ICP-MS

Plants were dried in an oven at 110 °C overnight, ground, and mineralized with concentrated nitric acid and 30% hydrogen peroxide for one hour each in a DigiPrep MS (SCP Science). The sample volume was adjusted to 25 mL, adding 10,000 ppb of Indium internal standard solution (SCP Science) for a final concentration of 50 pp and 2% (v/v) HNO3. Sample precipitates were filtered using a manifold (SCP Science) and 1.0 μm Digi filters (SCP Science). Dy concentrations were determined using an iCAP RQ quadrupole-based ICP-MS (Thermo Scientific). A certified reference (BCR-670, *Lemna minor*) was processed for quality control [32].

### 8. Dy detections with Deep UV LED light

*P. acinosa* leaves were incubated in 100 mM DyCl_3_, 100 mM CeCl_3,_ or water for 3 hours at room temperature. Then, small holes were generated with a thin needle (0.45 mm x 13 mm), and 3 μL of 2 M Na_2_WO_4_ were added to the pierced area, followed by a 5-minute incubation at room temperature to promote tungstate penetration through the epidermis. Following the Dy and tungstate treatment, the leaves were exposed to LED light (ThemoLabs, M275L4) at 275 nm and generated images using a smartphone camera under dark conditions

## Results

### 1. Fluorescence spectroscopy detects Dysprosium at 0.3 μM and higher concentrations

We first scanned the Dy absorbance using 100 mM DyCl_3_ in solution to determine the maximum Dy absorbance wavelengths. A 1 nm resolution scan from 250 to 500 nm identified seven major and five minor peaks (Supplemental Figure 1A). No additional peaks were detected between 500 to 700 nm wavelength region at 10 nm resolution (Supplemental Figure 1B). The maximum absorbance was 350 nm (Supplemental Figure 1A and 1B), consistent with the Dy UV-Vis absorption spectrum [29].

**Figure 1.**
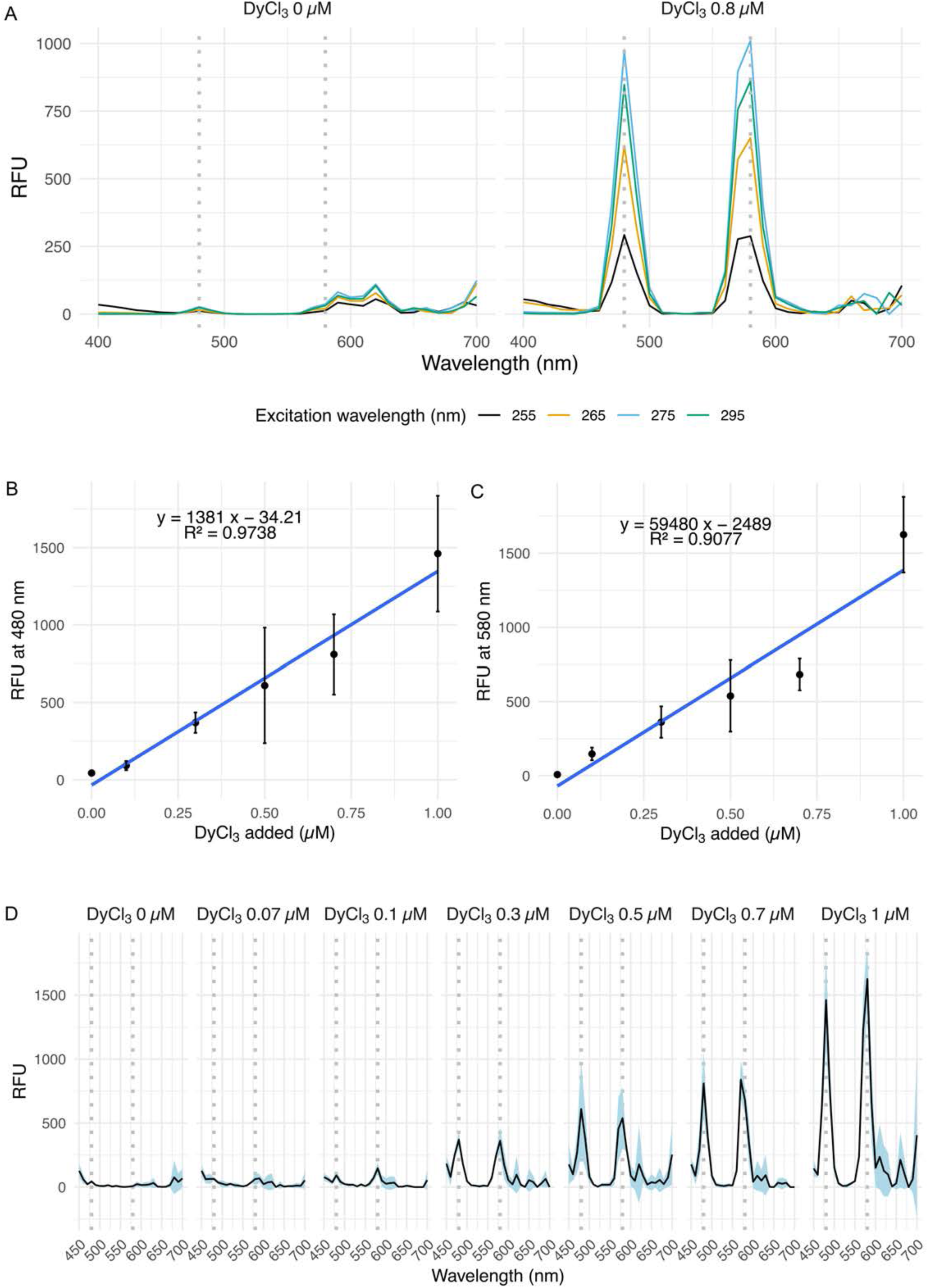
Dy emission signals and emission of minimum Dy levels detected by fluorescence spectroscopy in an aqueous DyCl3 solution. (A) Dy emission spectrum from 0 and 0.8 µM DyCl_3_ treated with 0.25 M Na_2_WO_4_, using 255, 265, 275, and 295 nm excitation wavelengths, 100 gain, and 60 µs time-resolved fluorescence. Lines indicate the mean measured value from 4 to 5 technical replicates. (B and C) Scatter plot and R-squared values between the 0-1 µM concentrations of DyCl_3_ with 0.25 M Na_2_WO_4_ and Dy emission at (B) 480 nm and (C) 580 nm measured with an excitation wavelength of 275 nm, 150 gain, and 60 µs time-resolved fluorescence. Data are mean ± SD from four replicates. (D) Dy emission spectra from 0-1 µM DyCl_3_ solution and 0.25 M Na_2_WO_4_ at 275 nm excitation, 150 gain, and 60 µs time-resolved fluorescence. The vertical dashed lines indicate 480- and 580-nm emission wavelengths. The blue-shaded ribbon indicates the standard deviation from three technical replicates.

Sodium tungstate (Na_2_WO_4_) enhances Dy fluorescence via excitation energy transfer [33,34]. We added 0.25 M Na_2_WO_4_ to DyCl_3_ solutions and evaluated the emission peaks at the excitation wavelengths identified in the absorbance assays. The characteristic Dy emission peaks were at 480 and 580 nm [30] (Figure 1A). We used time-resolved fluorescence in these assays because endogenous plant molecules cause autofluorescence [35], which can interfere with fluorescence spectra. Excitation between 255 and 295 nm generated robust Dy emission peaks, while 325 and 450 nm showed minimal emission (Figure 1, Supplemental Figure 1C and 1D). The highest fluorescence occurred with 275 nm excitation, which was selected for further experiments (Figure 1A, Supplemental Figure 1C and 1D).

A standard curve of DyCl_3_ concentrations (0 to 1 μM) with 0.25 M Na_2_WO_4_ and fluorescence emission at 480 and 580 nm showed a linear relationship (480 nm, R^2^ = 0.97) across the concentrations tested (Figure 1B - 1D). Dy signal was detectable from 0.1 μM DyCl_3_, with a robust and linear response from 0.3 μM in DyCl_3_ solutions.

### 2. Fluorescence lifetime measurements mitigate the contaminating effect of the plant matrix on Dy detection by fluorescence spectroscopy

We tested to see if Dy fluorescence was detectable in plant tissues. Homogenized leaf and stem tissue from *Phytolacca acinosa*, a species known to preferentially accumulate Dy [22,23], was mixed with 5 and 500 μM DyCl_3_ and 0.25 M Na_2_WO_4_. Dy was detectable at both concentrations (Figure 2A), but high autofluorescence was observed in the control plant tissues without Dy, particularly at 480 nm. The autofluorescence from endogenous plant molecules [35] confounded the Dy signal at 480 nm and made detection at the 5 μM concentration challenging. Implementing time-resolved fluorescence removed the background signal (Figure 2B). Since Dy has a microsecond-range fluorescence lifetime [36], delays of 20 to 80 μs after excitation substantially reduced the background signal from the plant matrix while still allowing Dy detection (Supplemental Figure 2).

**Figure 2.**
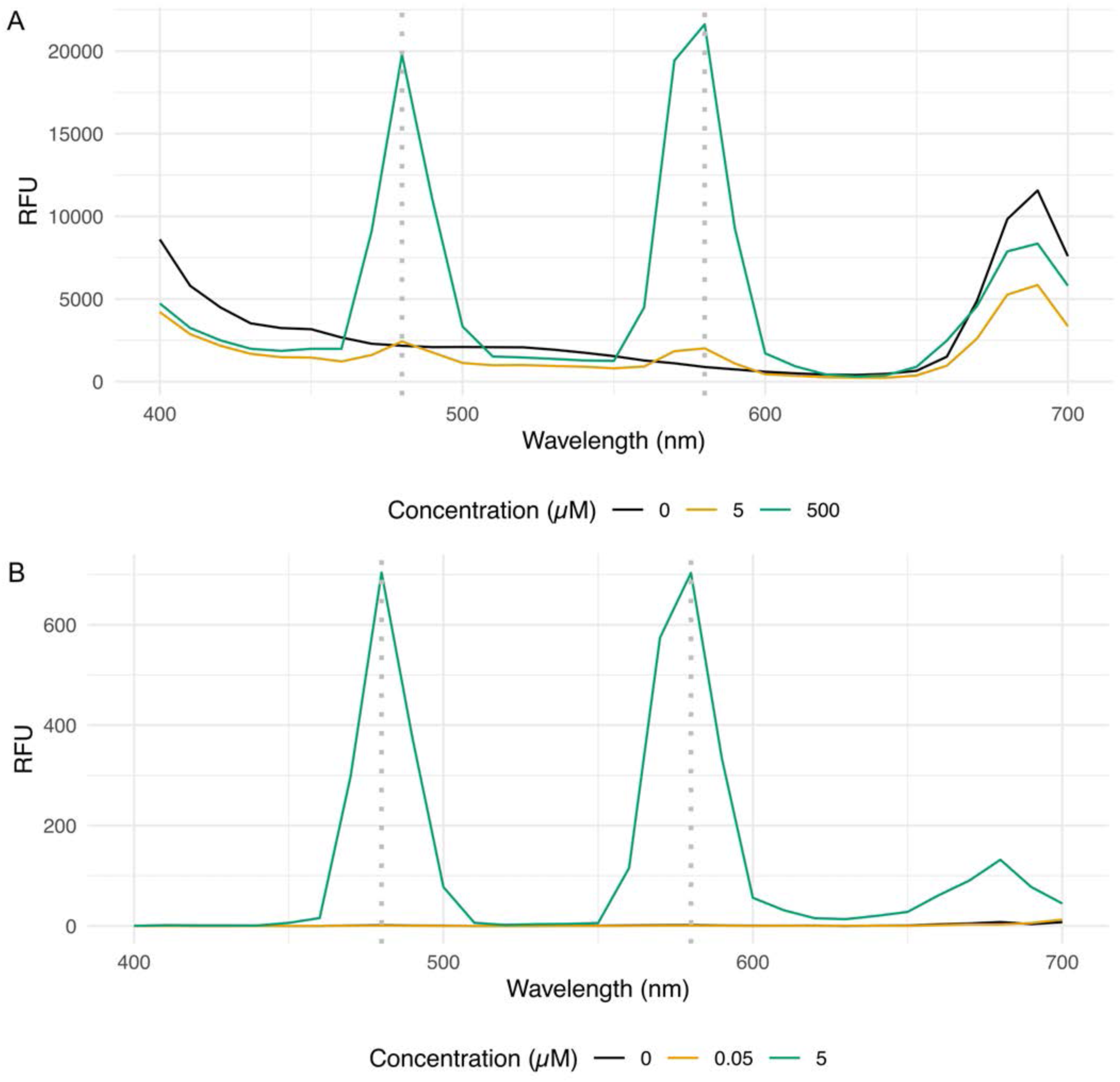
Dy screening in plant matrix with time-resolved fluorescence. (A and B) Dy emission spectra of 0, 5, and 500 µM DyCl_3_ with 0.25 M Na_2_WO_4_ in plant matrix (A) without and (B) with 60 µs time-resolved fluorescence and 100 gain. Lines indicate the mean measured value from five technical replicates.

We tested DyCl_3_ concentrations (0.05 - 25.6 µM) mixed with homogenized *P. acinosa* tissue and 0.25 M Na_2_WO_4_ to evaluate if Dy fluorescence is quantitative. We could detect the canonical Dy signal in the plant matrix background at concentrations above 0.8 µM (Supplemental Figure 3). The relative fluorescence units (RFU) at 480 nm and 580 nm after 275 nm excitation responded linearly to the Dy concentration (R^2^ > 0.99) (Figure 3A and 3B). ICP-MS results from the same sample confirmed these fluorescence measurements, with the fluorescence signal correlating with the ICP-MS determined concentrations (R^2^ > 0.99) (Figure 3C and 3D). Thus, Dy can be quantitatively detected at concentrations as low as 0.8 µM in a plant tissue background using fluorescence spectroscopy.

**Figure 3.**
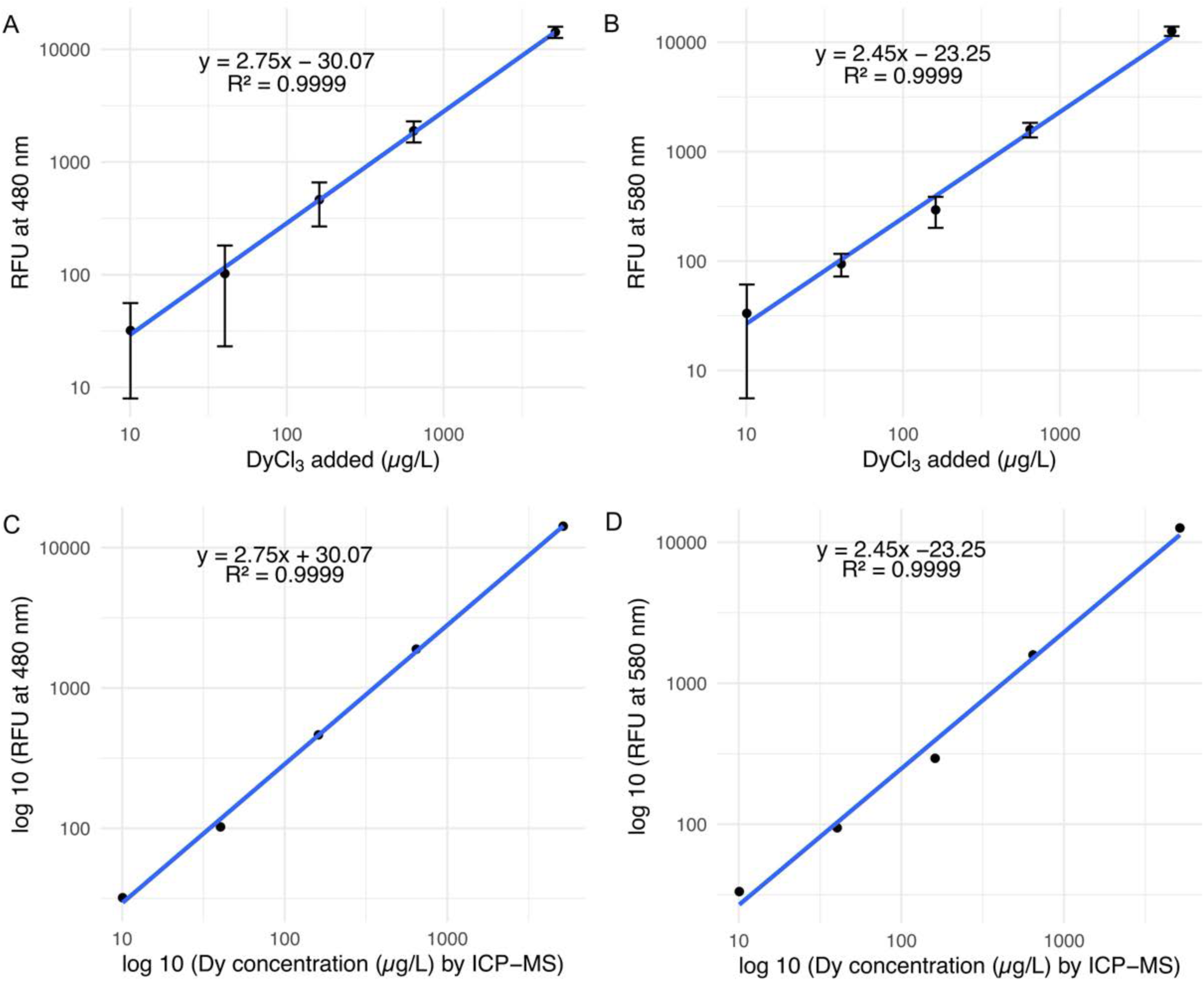
Dy detection in plant matrix by fluorescence spectroscopy. (A and B) Scatter plot of the concentrations of DyCl_3_ added to the plant matrix and the Dy emission at (A) 480 nm and (B) 580 nm. (C and D) Scatter plot of Dy concentrations measured by ICP-MS and Dy emission at (C) 480 nm and (D) 580 nm. Measurements were taken with an excitation wavelength of 275 nm, 150 gain, and 60 µs time-resolved fluorescence. Error bars indicate the standard deviation from four to five technical replicates.

### 3. Fluorescence spectroscopy can screen Dysprosium uptake from plant leaf tissue

We used time-resolved fluorescence to detect Dy in the leaves of plants grown in Dy solutions (Figure 4). *P. acinosa* plants were grown in modified Hoagland’s solution supplemented with 0, 5, 10, 15, and 20 mM DyCl_3_ for seven days. Leaf discs (2 mm) were taken from the lamina and midrib, treated with 0.5 M Na_2_WO_4_, and excited at 275 nm. The emission spectrum (450 to 700 nm) showed Dy signals in leaves treated with 5mM DyCl_3_ and higher but not in control plants (Supplemental Figure 4). Dy fluorescence at 580 nm increased with the Dy concentrations provided to the plants (Figure 4), indicating the method can screen for Dy translocated from roots to the leaves.

**Figure 4.**
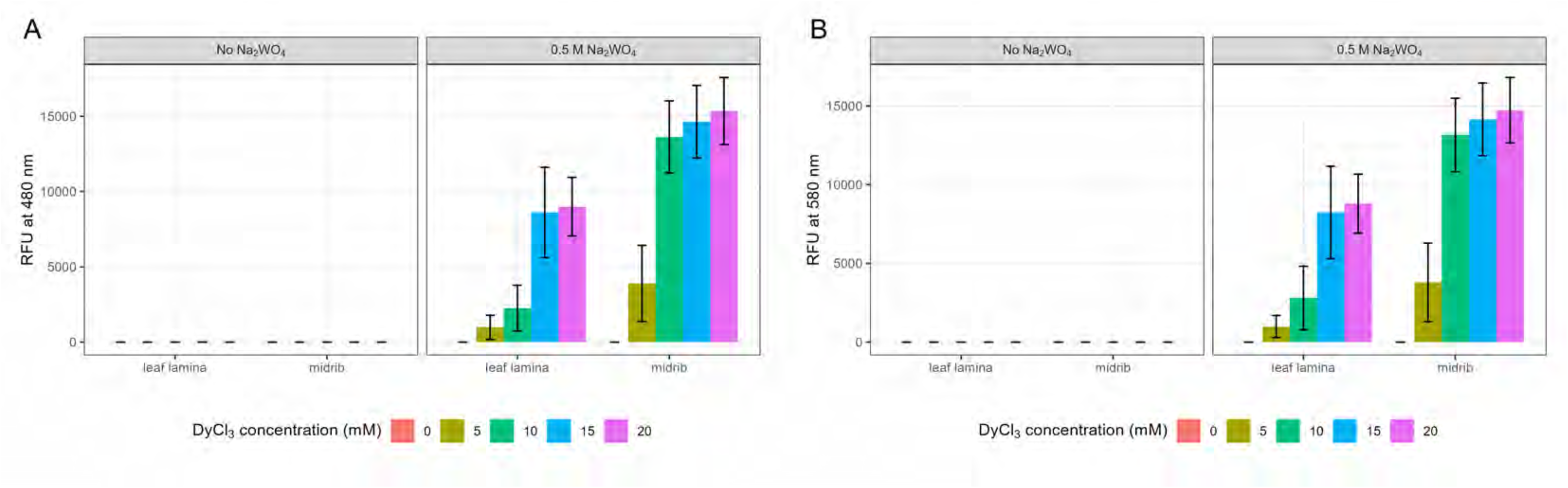
Dy screening of REE accumulation in plants from leaf discs. (A and B) Dy fluorescence at (A) 480 and (B) 580 nm from the leaf discs of *P. acinosa* plants grown in 0.25x Hoagland solution (without KH_2_PO_4_) supplemented with 0, 5, 10, 15, and 20 mM DyCl_3_ for seven days. Two discs, one from leaf lamina and another from midrib, were cut from four different plants and treated with 0.5 M Na_2_WO_4_. Dy fluorescence was detected from the leaf discs with a 275-nm excitation wavelength, 100 gain, and 60 µs time-resolved fluorescence. Data are mean ± the standard deviation.

### 4. Fluorescence spectroscopy can evaluate Dysprosium levels in living plants

We designed a method to detect Dy fluorescence directly from intact plant leaves in the field (Figure 5). This portable method does not implement time-resolved fluorescence, so plant autofluorescence is included, which is a limitation of this implementation. *P. acinosa* plants were treated with 10 and 50 mM DyCl_3_ or water (control) for two days. Using 248 nm excitation, Dy emissions increased with higher Dy treatments (Figure 5A). Autofluorescence was detected in control plants, especially at 480nm (Supplemental Figure 5A), but plants treated with DyCl_3_ were distinguishable from controls and from each other (10 vs. 50 mM) (Figure 5A and Supplemental Figure 5A).

**Figure 5.**
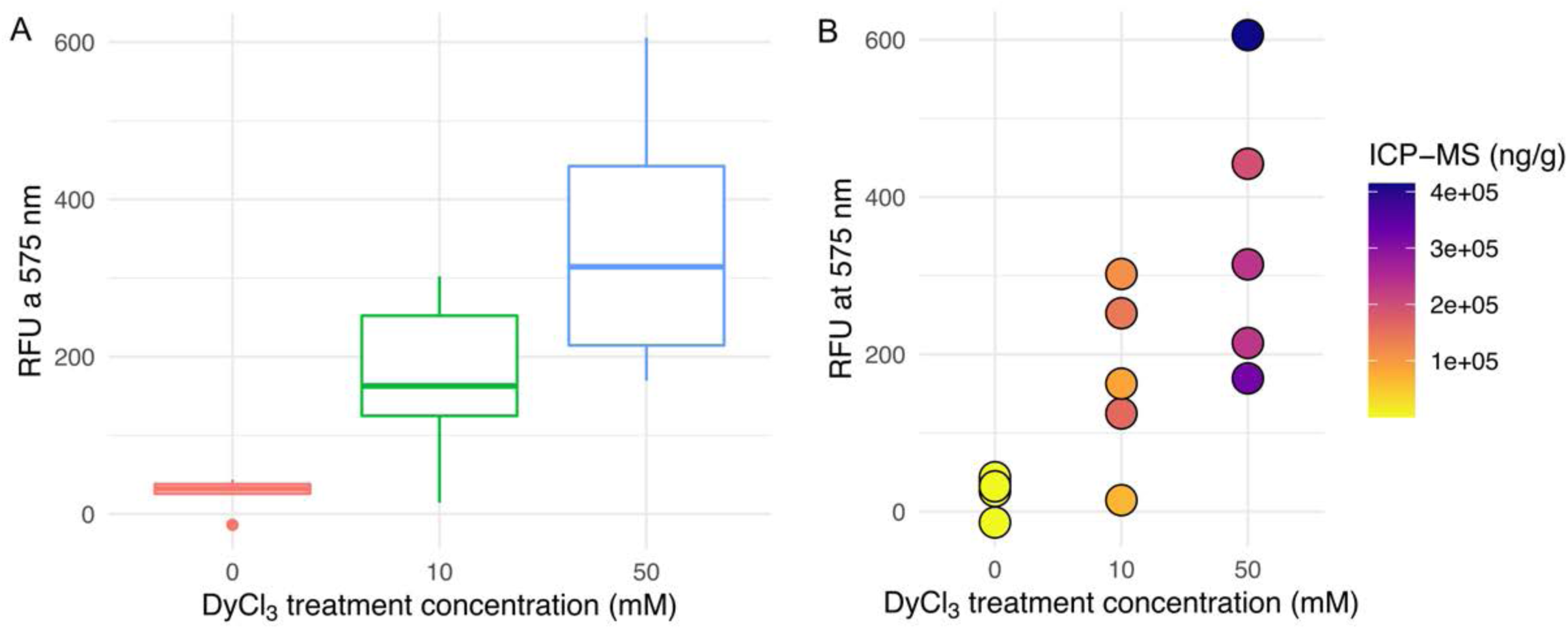
Non-destructive screening of Dy in plant tissue. (A) Box and whisker plots of Dy emission at 575 nm wavelength after excitation at 248 nm in the tissue of plants treated with DyCl_3_. Each box represents a measurement from 5 biological replicates (individual plants). (B) Point plot of Dy emission at 575 nm wavelength after 248 nm excitation and Dy concentrations measured by ICP-MS represented by a color gradient. Yellow-hued colors represent lower concentration values as determined by ICP-MS, and blue represents high concentration values. Each point represents an individual plant.

After the fluorescence measurement, leaves were analyzed by ICP-MS. Both Dy emission and ICP-MS measurements showed increased Dy detection variance with higher treatment concentrations (Figure 5B, Supplemental Figure 5B, and 5C). A positive correlation was found between Dy emission and ICP-MS measurements (Figure 5B and Supplemental Figure 5C). Discrepancies in some individual samples could be due to the location assayed and sampling protocols. The fluorescence measures a specific spot, while ICP-MS measures the combined Dy concentration of several leaves.

Variations in the plant autofluorescence also contribute to differences in the background signal. Developing a portable device with delayed fluorescence would reduce this background noise. Although further improvements to portable detection will increase the accuracy, this system offers a rapid first-pass screening to identify plants with high Dy uptake levels.

### 5. Detection of Dysprosium at a Standoff Distance

The Dy fluorescence signal in the presence of Na_2_WO_4_ is bright and visible to the eye. Therefore, since imaging at a distance could increase the high-throughput screening, we tested the ability to detect the Dy uptake in plants at a standoff distance. We treated *P. acinosa* leaves with 100 mM DyCl_3_ for 3 hours and induced tissue penetration of Na_2_WO_4_ by piercing the leaves several times with a thin needle (0.45 mm x 13 mm) and adding 3 µL of 2 M Na_2_WO_4_ on the pierced area. We illuminated the leaf with an LED of 275 nm and took a picture with a smartphone. Control samples included leaves treated with water (+/- Na_2_WO_4_) or cerium (+/- Na_2_WO_4_). The Dy fluorescence is already evident at 3h in plants exposed to this high Dy concentration from the smartphone pictures of the leaf areas treated with Dy and Na_2_WO_4_ (Figure 6). There is no discernable fluorescence from the controls with cerium, water, or Dy without Na_2_WO_4_. This indicates that Dy detection can be visualized at a standoff distance, which could facilitate rapid, high-throughput screening by remote monitoring.

**Figure 6.**
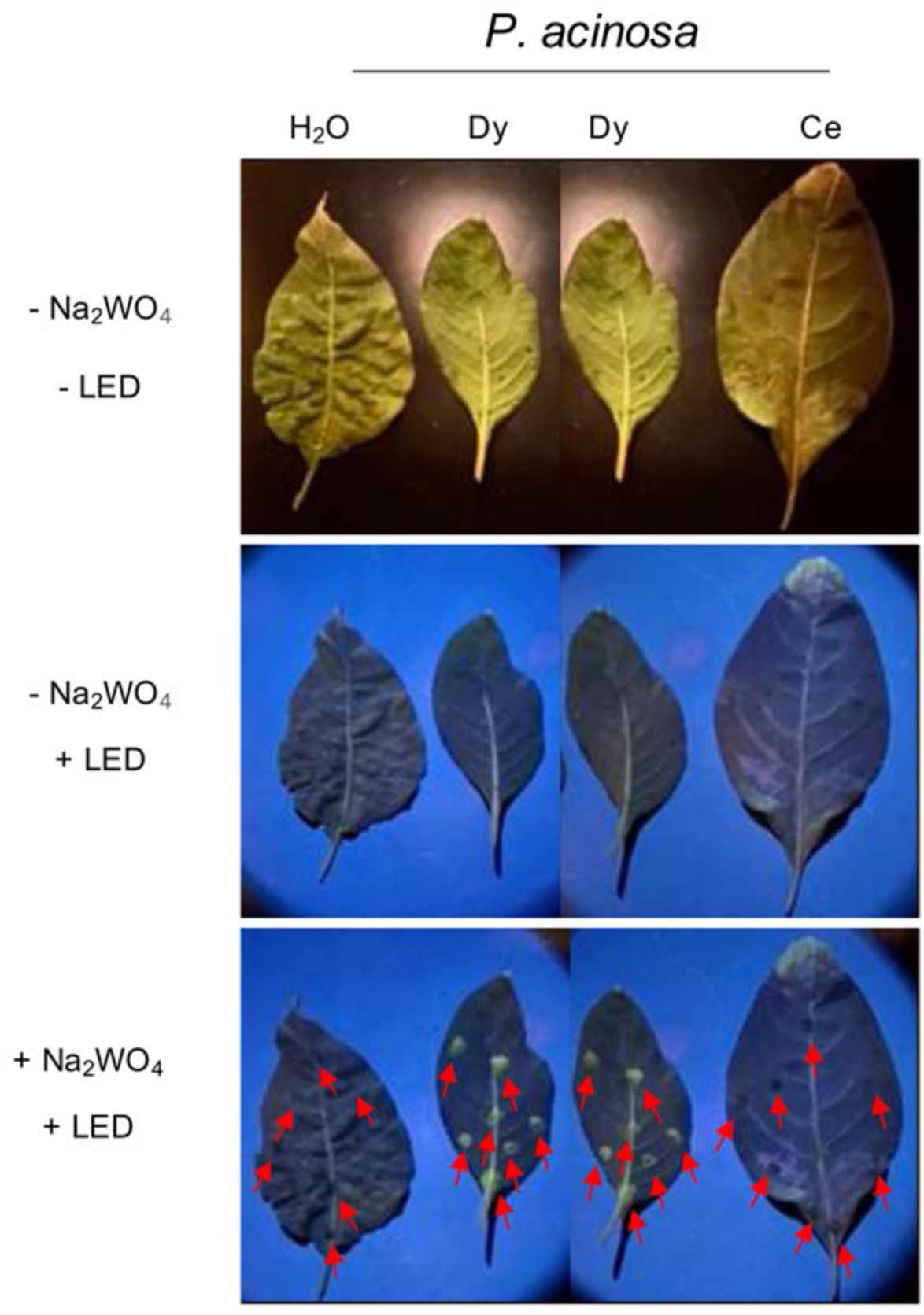
Standoff detection of Dy uptake in plant leaves. Fluorescence from *P. acinosa* leaves incubated with water (H_2_O), Ce, and Dy, captured by a smartphone camera. Image panels are: Top panel, - Na_2_WO_4_, - LED: an image of punctured leaves before Na_2_WO_4_ treatment and 275 nm LED exposure. Middle panel, - Na_2_WO_4_, + LED: an image of punctured leaves, before Na_2_WO_4_ treatment exposed to the 275 nm LED light. Bottom panel, + Na_2_WO_4_, + LED: an image of fluorescence from punctured leaves treated with Na_2_WO_4_ exposed to 275 nm LED light. Arrows indicate where Na_2_WO_4_ treatments were placed.

## Discussion

Phytomining is a promising technique for enhancing the recovery of REEs while minimizing the environmental footprint associated with traditional mining methods. The number of species identified as suitable for metal accumulation is limited. For example, Liu et al. identified only 22 plant species as REE hyperaccumulators or potential REE hyperaccumulators [37]. To advance REE phytomining, it is crucial to identify plants that can absorb unusually high REE concentrations and to understand how environmental factors and microbial interactions influence REE uptake. Strategies to improve plant uptake through genetic modification, microbial treatments, or other environmental factors will further enhance phytomining efforts. Critically, the selection of improved genotypes and environmental parameters that improve REE uptake and specificity is limited by the need for rapid, high-throughput screening methods. Rapid screening methods to identify REE uptake will accelerate the assessment of natural and engineered genotypes and enhance our understanding of how external factors affect REE uptake by plants.

Fluorescence spectroscopy is a fast method for quantifying Dy levels in plant tissue. Using a plate reader for these assays offers the advantage of time-resolved fluorescence, which overcomes the challenges of the autofluorescence produced by the plant. Dy produces a longer- lived luminescence than the compounds responsible for plant autofluorescence (Figure 2) [38]. This method can differentiate Dy levels in various parts of the leaf, such as the lamina and the midrib, providing insights into the localization and distribution of Dy within different plant organs (Figure 4). This screening can be done with only a small tissue sample provided by a 2 mm diameter leaf punch. This approach enables rapid screening of Dy emission in the tissue while preserving the rest of the plant’s aerial parts.

Our UV fluorescence approach also enables rapid, non-destructive detection of Dy in plant tissue in field settings (Figure 5). This non-destructive sampling can facilitate monitoring the same plant across time, allowing insights into dynamics and how different environmental treatments affect Dy uptake. Non-destructive Dy monitoring can facilitate tracking Dy levels at different growth stages and how Dy is distributed across different tissues as the plant matures. Time-course monitoring of individual plants can improve the accuracy of studies examining how Dy uptake is affected by environmental changes, such as the addition of microbes or alterations to the soil environment. These findings can aid in optimizing growth conditions to enhance Dy uptake. Additionally, continuous Dy monitoring can help determine the optimal harvesting time to maximize Dy recovery, ensuring plants are collected when Dy concentrations peak.

The present instrument we developed for field-based assays lacks time-resolved fluorescence. Therefore, the plant’s autofluorescence can interfere with the Dy signal. It is crucial to account for autofluorescence when comparing plant species, as different plants have varying background signals that can change in response to environmental changes [35]. Although currently limited by the lack of time-resolved fluorescence, this field-based evaluation can be used as an initial screen. Leaf discs collected from the top Dy accumulation candidates identified by this screen can be evaluated using time-resolved fluorescence in the laboratory. Developing a portable device with time-resolved capability would enhance sensitivity and accuracy for non-destructive Dy detection in the field and herbarium samples, similar to advancements achieved with XRF [27].

Additionally, the emission from Dy complexed with Na_2_WO_4_ is visible to the naked eye (Figure 6), which opens up practical applications for plant analysis. Standoff detection techniques can be employed using a phenotyping device from a distance to measure plant Dy plant uptake after treating the plants with Na_2_WO_4_. This method is particularly useful for detecting plants that accumulate high concentrations of Dy, as the visible emission will highlight those plants more effectively. This approach would simplify and improve the efficiency of plant screening to identify high-Dy accumulating plants in research and agricultural settings.

This investigation focused on developing a quick method to screen Dy in plant tissue. The plants grew in high Dy concentrations that were applied hydroponically in the form of DyCl_3_. This system allowed us to focus on Dy detection and eliminate variability in uptake due to environmental variables. However, this is an artificial delivery of REEs [15]. This delivery system demonstrates that the detection correlates with measures by ICP-MS and can identify plants with uptake from those without. The ability to measure down to 1 μM in the plant background indicates that this detection method can screen Dy in plants and potentially identify new Dy hyperaccumulators.

Fluorescence assays can be expanded to screen for other REEs in plant tissue. REEs exhibit a wide range of emission characteristics across the UV-Vis-NIR spectrum, which makes them detectable using fluorescence spectroscopy. This expansion offers new opportunities for research into REE uptake by plants and for advancing phytomining techniques.

## Conclusion

In conclusion, this work demonstrates the development of rapid and non-destructive methods for detecting Dy in plant tissues. The ability to quantitatively assess Dy levels in living plant tissues represents a significant advancement in phytomining. The use of fluorescence spectroscopy offers promising avenues for improving the efficiency and effectiveness of phytomining research. By quickly identifying plants with high REE uptake, these methods can accelerate the screening and selection of potential hyperaccumulators, facilitating further research and optimization of phytomining practices. Evaluating the response of the same plant across time will inform how environmental factors affect Dy uptake. The unique spectral characteristics of other REEs suggest that this method could be adapted to quantify the uptake of several different REEs. Expanding these assays to include other REEs could greatly enhance our understanding of plant-REE interactions and contribute to more sustainable and environmentally friendly mining solutions. In summary, adapting fluorescence spectroscopy for REE detection can enhance phytomining research by enabling the rapid quantification and localization of REEs, paving the way for more sustainable and efficient resource recovery practices.

## Abbreviations

Ce: Cerium
Dy: Dysprosium
DyCl_3_: Dysprosium chloride
La: lanthanum
ICP-AES: ICP atomic emission spectroscopy
ICP-MS: Inductively Coupled Plasma Mass Spectrometry
ICP-OES: ICP optical emission spectroscopy
Nd: neodymium
REE: Rare Earth Element
Na_2_WO_4_: Sodium tungstate
UV: ultraviolet
UV-Vis-NIR: ultraviolet-visible-near-infrared
XRF: X-ray fluorescence
Y: yttrium.

## Declarations

### Ethics, Consent to Participate, and Consent to Publish

Not applicable.

### Data Availability Statement

All data generated or analyzed during this study are included in this published article (and its Supplementary Information files).

### Authors’ Contributions

M.W.K. and C.J.D. Designed the initial project. E.H.P., A.T.H., and K.L. optimized the detection assays. E.H.P, A.T.H., A.N.H, C.R., D.B., and M.W.K. performed the measurements.

M.W.K. developed the imaging system used for Figures 5 and 6. E.H.P, M.W.K., and C.J.D. prepared the initial draft. All authors contributed to the writing and review of the manuscript.

## Acknowledgements

We would like to thank Kendrick Turner, Blake Bextine for assay development advice and Liz Rylott for guidance and advice on the manuscript preparation.

---

**Supplemental Figure 1. Dy absorbance spectra and emission signals.** (A) Dy absorbance spectra scanned from 250 - 500 nm at 1-nm intervals for 100 mM DyCl_3_. (B) Dy absorbance spectra from 100 mM DyCl_3_ in aqueous solution scanned from 250 - 700 nm at 10-nm intervals. Lines indicate the mean measured value from four to five technical replicates. (C and D) Dy emission from 255, 265, 275, 295, 325, 350, 365, 390, 425, and 450 nm excitation wavelengths from 0 and 0.8 µM DyCl_3_ in aqueous solutions with 0.25 M Na_2_WO_4_. Emission was measured at (B) 480 and (D) 580 nm using a 100 gain and 60 µs time-resolved fluorescence. Error bars indicate the standard deviation from four to five technical replicates.

**Supplemental Figure 2. Dy screening in plant matrix with time-resolved fluorescence.** Time-resolved fluorescence lengths of 0, 20, 30, 40, 50, and 60 µs were tested to evaluate the detection of Dy in the presence of *P. acinosa* tissue. A solution of 3.2 µM Dy and 0.5 M Na_2_WO_4_ was mixed with *P. acinosa* homogenized leaf and stem tissue. Dy fluorescence was measured with a 275-nm excitation wavelength and 150 gain. Lines indicate the mean measured value from five technical replicates.

**Supplemental Figure 3. Dy emission in plant matrix.** Dy emission spectra of 0.05 to 25.6 µM DyCl_3_ concentrations with 0.5 M Na_2_WO_4_ in plant matrix. Measurements were taken with an excitation wavelength of 275 nm and 150 gain with 60 µs time-resolved fluorescence. The blue-shaded ribbon indicates the standard deviation from four to five technical replicates.

**Supplemental Figure 4. Dy emission spectra of REE uptake in plants from leaf discs.** Leaf discs (2 mm) were taken from *P. acinosa* plants grown in 0.25x modified Hoagland solution supplemented with 0, 5, 10, 15, and 20 mM DyCl_3_ for seven days. Two discs, one from leaf lamina and another from midrib, were cut from four different plants and treated with 0.5 M Na_2_WO_4_. Dy fluorescence was measured from the leaf discs with a 275 nm excitation wavelength and 100 gain with 60 µs time-resolved fluorescence. The vertical dashed lines indicate 480 and 580 nm emission wavelengths.

**Supplemental Figure 5. Non-destructive Dy detection in plant tissue.** (A) Box and whisker plots of Dy emission at 480 nm wavelength after excitation at 248 nm in the tissue of plants treated with DyCl_3_. (B) Box and whisker plots of Dy concentrations in plants treated with DyCl_3_ measured by ICP-MS. (C) Point plot of Dy emission at 480 nm wavelength after 248 nm excitation and Dy concentrations measured by ICP-MS represented by a color gradient. Yellow-hued colors represent lower concentration values as determined by ICP-MS, and blue represents high concentration values. Each box represents a measurement from 5 biological replicates. Each point represents an individual plant.

## REFERENCES

1. Voncken JHL. The rare earth elements. 1st ed. Cham, Switzerland: Springer International Publishing; 2016.

2. Seo Y, Morimoto S. Comparison of dysprosium security strategies in Japan for 2010–2030. Resour Policy. 2014;39:15–20.

3. Xiao S, Geng Y, Pan H, Gao Z, Yao T. Uncovering the key features of dysprosium flows and stocks in China. Environ Sci Technol. 2022;56:8682–90.

4. Eheliyagoda D, Ramanujan D, Veluri B, Liu Q, Liu G. Tracing the multiregional evolution of the global dysprosium demand-supply chain. Resour Conserv Recycl. 2023;199:107245.

5. Riaño S, Binnemans K. Extraction and separation of neodymium and dysprosium from used NdFeB magnets: an application of ionic liquids in solvent extraction towards the recycling of magnets. Green Chem. 2015;17:2931–42.

6. Balaram V. Rare earth elements: A review of applications, occurrence, exploration, analysis, recycling, and environmental impact. Geoscience Frontiers. 2019;10:1285–303.

7. Dutta T, Kim K-H, Uchimiya M, Kwon EE, Jeon B-H, Deep A, et al. Global demand for rare earth resources and strategies for green mining. Environ Res. 2016;150:182–90.

8. Li X-Y, Ge J-P, Chen W-Q, Wang P. Scenarios of rare earth elements demand driven by automotive electrification in China: 2018–2030. Resour Conserv Recycl. 2019;145:322–31.

9. Talan D, Huang Q. A review of environmental aspect of rare earth element extraction processes and solution purification techniques. Miner Eng. 2022;179:107430.

10. Peiravi M, Dehghani F, Ackah L, Baharlouei A, Godbold J, Liu J, et al. A review of rare-earth elements extraction with emphasis on non-conventional sources: Coal and coal byproducts, iron ore tailings, apatite, and phosphate byproducts. Min Metall Explor. 2021;38:1–26.

11. Mwewa B, Tadie M, Ndlovu S, Simate GS, Matinde E. Recovery of rare earth elements from acid mine drainage: A review of the extraction methods. Journal of Environmental Chemical Engineering. 2022;10:107704.

12. Dinh T, Dobo Z, Kovacs H. Phytomining of rare earth elements – A review. Chemosphere. 2022;297:134259.

13. van der Ent A, Baker AJM, Echevarria G, Simonnot M-O, Morel JL. Agromining: Farming for metals: Extracting unconventional resources using plants. Springer Nature; 2020.

14. Yang X, Feng Y, He Z, Stoffella PJ. Molecular mechanisms of heavy metal hyperaccumulation and phytoremediation. J Trace Elem Med Biol. 2005;18:339–53.

15. van der Ent A, Rylott EL. Inventing hyperaccumulator plants: improving practice in phytoextraction research and terminology. Int J Phytoremediation. 2024;26:1379–82.

16. Rylott EL, Bruce NC. Plants to mine metals and remediate land. Science. 2022;377:1380–1.

17. Tognacchini A, Rosenkranz T, van der Ent A, Machinet GE, Echevarria G, Puschenreiter M. Nickel phytomining from industrial wastes: Growing nickel hyperaccumulator plants on galvanic sludges. J Environ Manage. 2020;254:109798.

18. Mesjasz-Przybylowicz J, Przybylowicz W, Barnabas A, van der Ent A. Extreme nickel hyperaccumulation in the vascular tracts of the tree Phyllanthus balgooyi from Borneo. New Phytol. 2016;209:1513–26.

19. Isnard S, L’Huillier L, Paul ALD, Munzinger J, Fogliani B, Echevarria G, et al. Novel Insights into the hyperaccumulation syndrome in Pycnandra (Sapotaceae). Front Plant Sci. 2020;11:559059.

20. Chaney R. Phytoextraction and phytomining of soil nickel. Nickel in soils and plants [Internet]. 2018; Available from: https://www.taylorfrancis.com/chapters/edit/10.1201/9781315154664-15/phytoextraction-phytomining-soil-nickel-rufus-chaney

21. Liu C, Liu W-S, van der Ent A, Morel JL, Zheng H-X, Wang G-B, et al. Simultaneous hyperaccumulation of rare earth elements, manganese and aluminum in Phytolacca americana in response to soil properties. Chemosphere. 2021;282:131096.

22. Grosjean N, Le Jean M, Berthelot C, Chalot M, Gross EM, Blaudez D. Accumulation and fractionation of rare earth elements are conserved traits in the Phytolacca genus. Sci Rep. 2019;9:18458.

23. Yuan M, Liu C, Liu W-S, Guo M-N, Morel JL, Huot H, et al. Accumulation and fractionation of rare earth elements (REEs) in the naturally grown Phytolacca americana L. in southern China. Int J Phytoremediation. 2018;20:415–23.

24. Zawisza B, Pytlakowska K, Feist B, Polowniak M, Kita A, Sitko R. Determination of rare earth elements by spectroscopic techniques: a review. J Anal At Spectrom. 2011;26:2373.

25. Huang CL, Schulte EE. Digestion of plant tissue for analysis by ICP emission spectroscopy. Commun Soil Sci Plant Anal. 1985;16:943–58.

26. Liu W-S, van der Ent A, Erskine PD, Morel JL, Echevarria G, Spiers KM, et al. Spatially resolved localization of lanthanum and cerium in the rare earth element hyperaccumulator fern Dicranopteris linearis from China. Environ Sci Technol. 2020;54:2287–94.

27. Goudard L, Blaudez D, Sirguey C, Purwadi I, Invernon V, Rouhan G, et al. Prospecting for rare earth element (hyper)accumulators in the Paris Herbarium using X-ray fluorescence spectroscopy reveals new distributional and taxon discoveries. Ann Bot. 2024;133:573–84.

28. Sato M, Kim SW, Shimomura Y, Hasegawa T, Toda K, Adachi G. Chapter 278 - Rare earth- doped phosphors for white light-emitting diodes. In: Jean-Claude B, Vitalij K. P, editors. Handbook on the Physics and Chemistry of Rare Earths. Elsevier; 2016. p. 1–128.

29. Runowski M, Stopikowska N, Lis S. UV-Vis-NIR absorption spectra of lanthanide oxides and fluorides. Dalton Trans. 2020;49:2129–37.

30. Sruthi P, Swapna K, Kumar JVS, Mahamuda S, Venkateswarulu M, Amer D, et al. Dysprosium concentration-dependent fluorescent properties of antimony lead Oxyfluoroborate glasses. Chem Phys Lett. 2022;787:139210.

31. Banah H, Balint-Kurti PJ, Houdinet G, Hawkes CV, Kudenov M. The quantification of southern corn leaf blight disease using deep UV fluorescence spectroscopy and autoencoder anomaly detection techniques. PLoS One. 2024;19:e0301779.

32. Berky AJ, Weinhouse C, Vissoci J, Rivera N, Ortiz EJ, Navio S, et al. In utero exposure to metals and birth outcomes in an artisanal and small-scale gold mining birth cohort in Madre de Dios, Peru. Environ Health Perspect. 2023;131:97008.

33. Alberti G, Massucci MA. Spectrofluorometric trace determination of trivalent samarium, europium, terbium, and dysprosium by sodium tungstate solution. Anal Chem. 1966;38:214–6.

34. Xu L-J, Xu G-T, Chen Z-N. Recent advances in lanthanide luminescence with metal-organic chromophores as sensitizers. Coord Chem Rev. 2014;273–274:47–62.

35. Donaldson L. Autofluorescence in plants. Molecules [Internet]. 2020;25. Available from: 10.3390/molecules25102393

36. Berezin MY, Achilefu S. Fluorescence lifetime measurements and biological imaging. Chem Rev. 2010;110:2641–84.

37. Liu C, Yuan M, Liu W-S, Guo M-N, Zheng H-X, Huot H, et al. Element case studies: Rare earth elements. In: van der Ent A, Baker AJM, Echevarria G, Simonnot M-O, Morel JL, editors. Agromining: Farming for metals: Extracting unconventional resources using plants. Cham: Springer International Publishing; 2021. p. 471–83.

38. Hagan AK, Zuchner T. Lanthanide-based time-resolved luminescence immunoassays. Anal Bioanal Chem. 2011;400:2847–64.

