## Supplementary figures and images for "Nondestructive Detection and Quantification of Dysprosium in Plant Tissues"

### Supplemental Figure 1

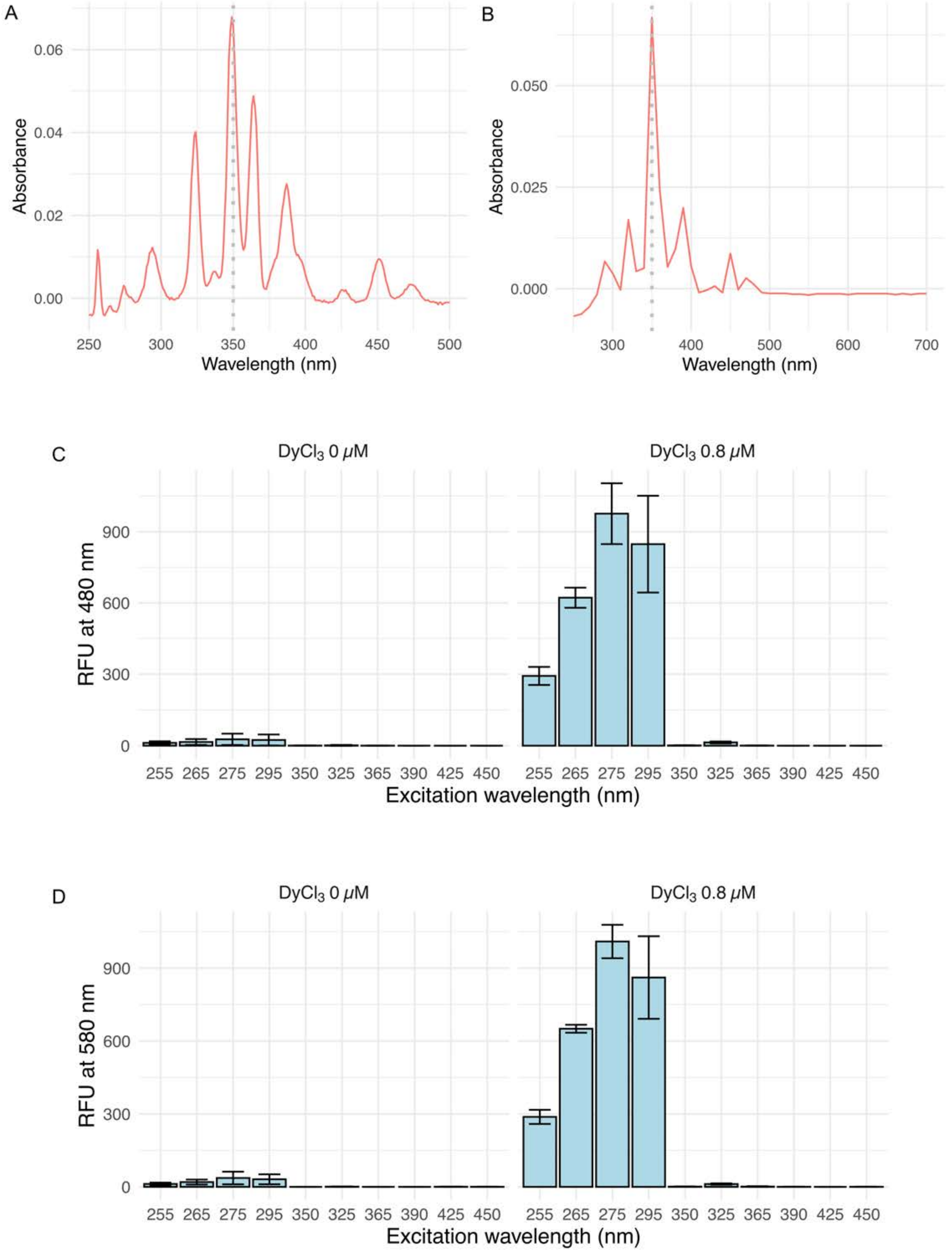

### Supplemental Figure 2

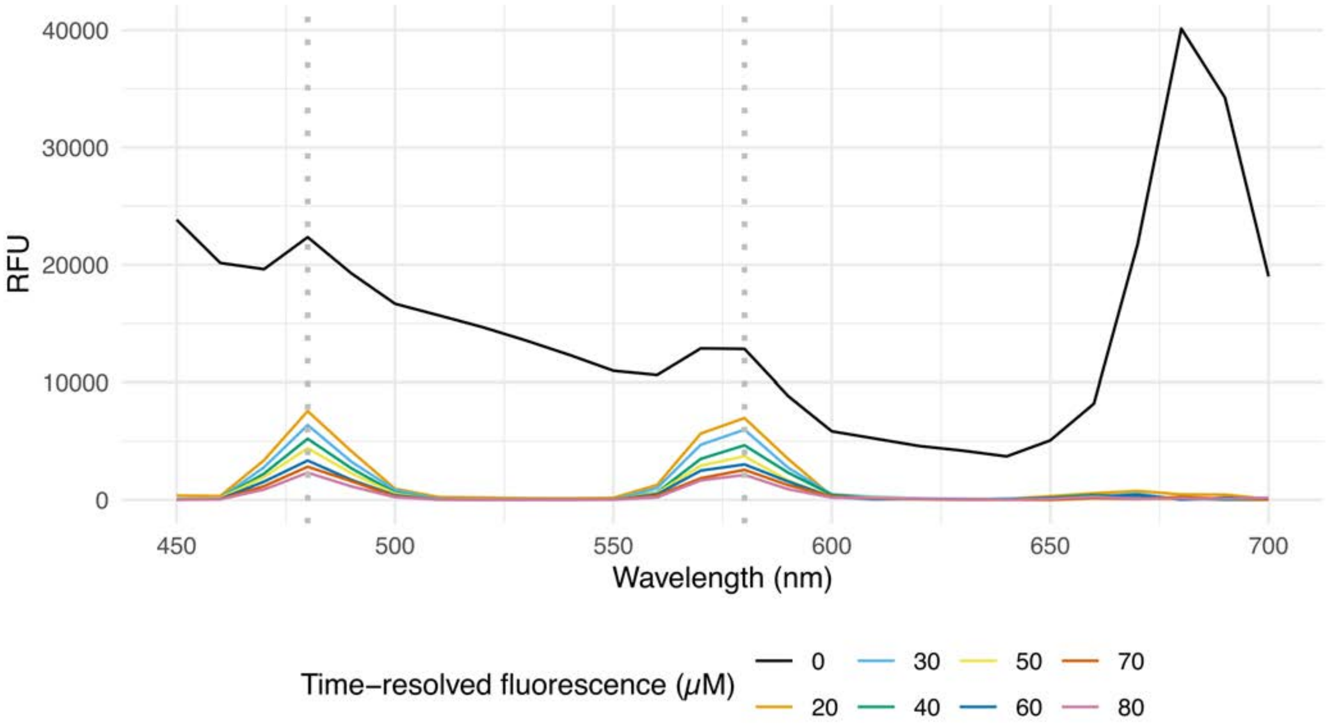

### Supplemental Figure 3

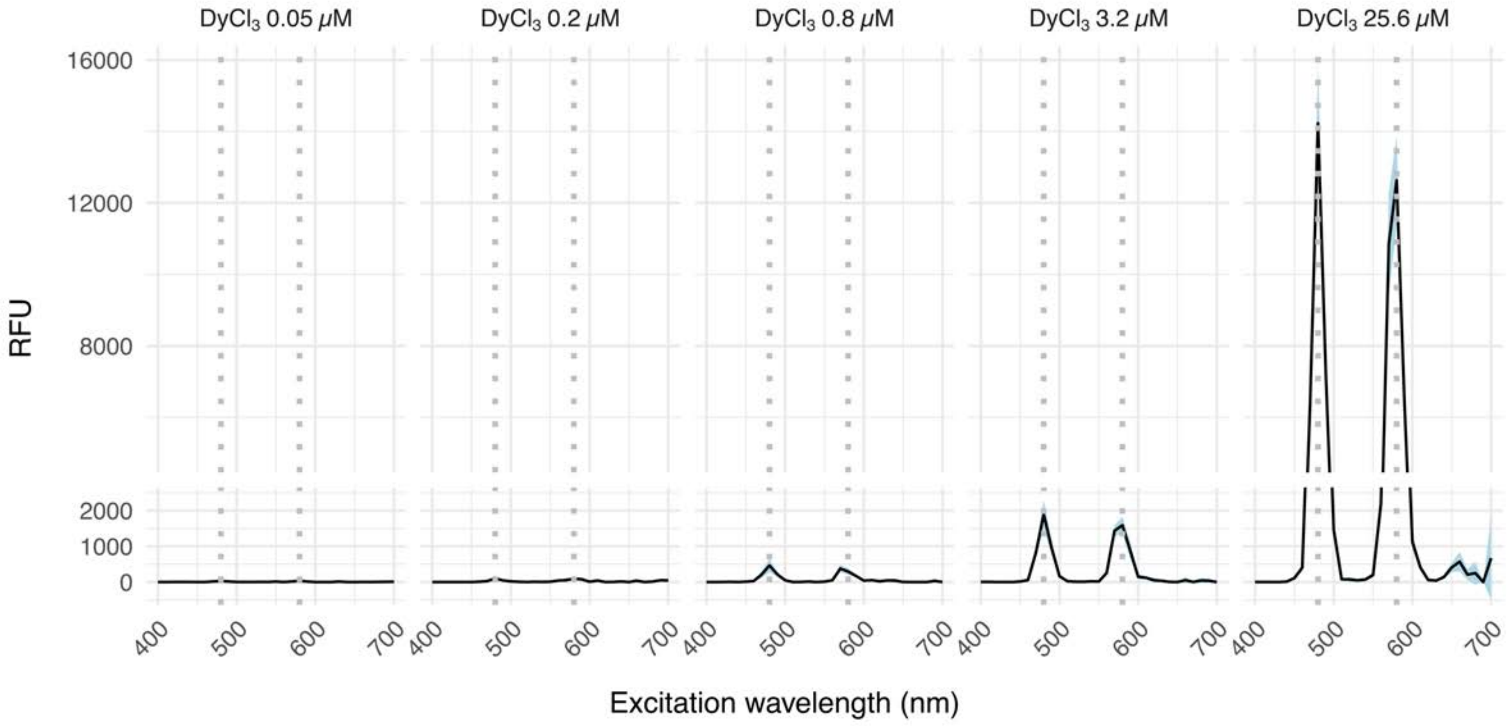

### Supplemental Figure 4

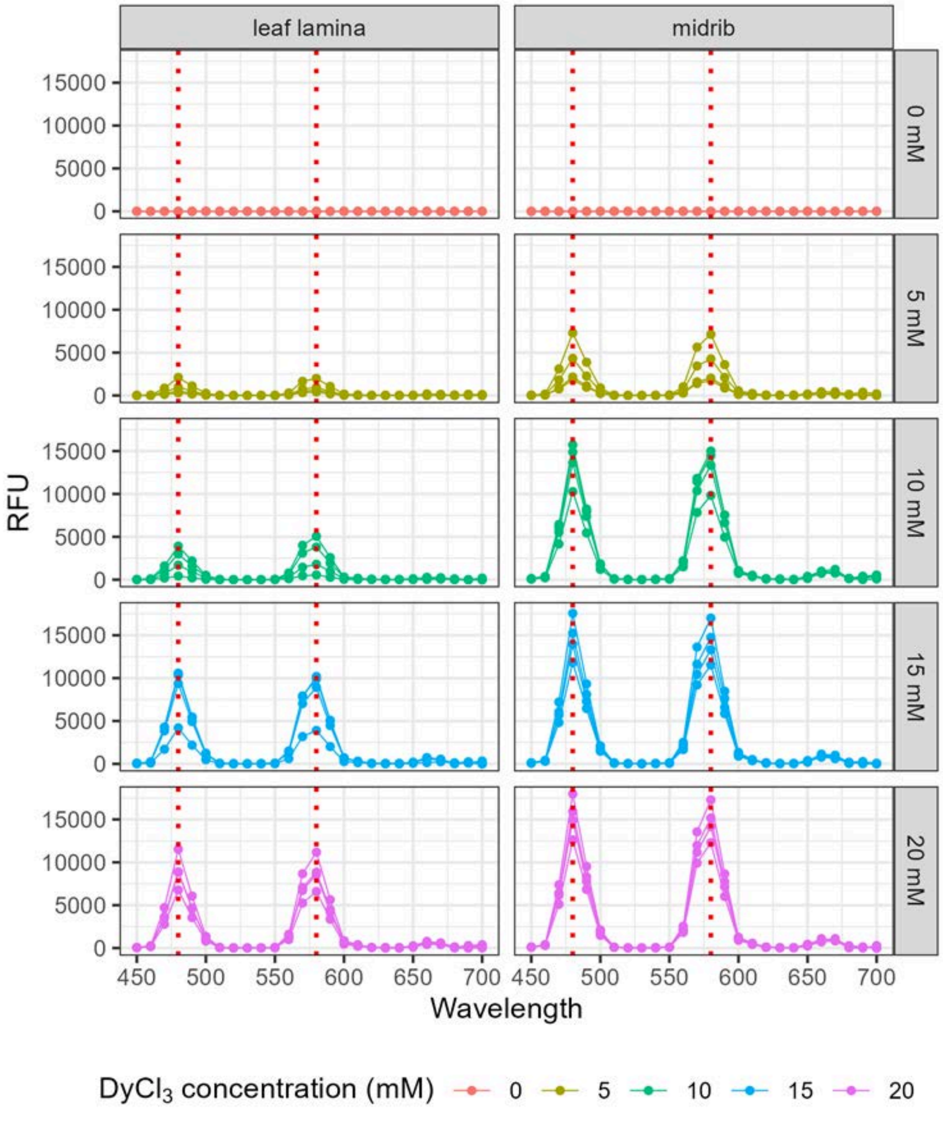

### Supplemental Figure 5

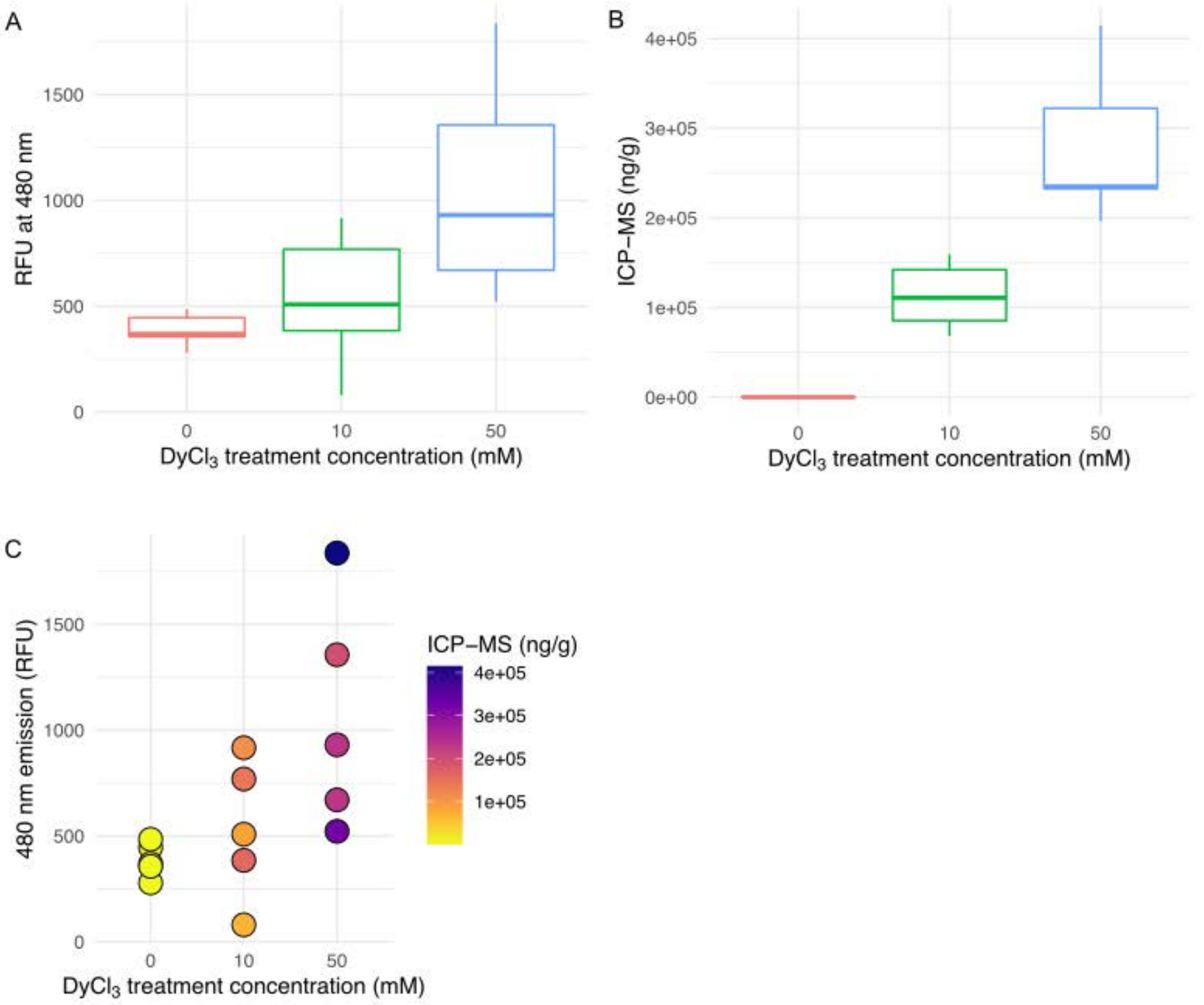
